# Diffusion of DNA on Atomically Flat 2D Material Surfaces

**DOI:** 10.1101/2023.11.01.565159

**Authors:** Dong Hoon Shin, Sung Hyun Kim, Kush Coshic, Kenji Watanabe, Takashi Taniguchi, Gerard Verbiest, Sabina Caneva, Aleksei Aksimentiev, Peter G. Steeneken, Chirlmin Joo

## Abstract

Accurate localization of biomolecules is pivotal for understanding biological processes. Utilizing the atomically flat surface of 2D materials offers a promising route to achieve this without the need for tethering or constraining. Here we comprehensively investigate the binding and diffusion of DNA on hexagonal boron nitride (hBN) surfaces. Our findings reveal non-specific binding of DNA to pristine hBN, with subsequent diffusion and confinement within the 2D plane. Through single-molecule experiments and computational techniques, we explore DNA dynamics, and the effects of defects, step edges and domain boundaries on the motion, which gives insights on the interactions between solid-state surfaces and biomolecules. By engineering a narrow hBN ribbon structure, we enhance confinement, demonstrating its potential in nanofluidic guiding of biomolecules. Our 2D platform serves as a proving ground for next generation high-throughput single-molecule manipulation techniques for enabling applications in biotechnology and nanotechnology.

## INTRODUCTION

The localization of biomolecules such as DNA and proteins is an indispensable endeavour, bearing immense significance in the comprehensive study of biological entities and their interactions^1–6^. Achieving this aim without the need for tethering or constraining these molecules would not only facilitate the accurate detection and identification of single biomolecules but also empower the direct monitoring and controlling of bio-chemical reactions and the detailed characterization of their properties^7–11^. A promising avenue involves characterizing the properties of biomolecules as they traverse a precisely engineered two-dimensional (2D) surface. This surface serves as a nanoscale 2D biocharacterization processing line, enabling the isolation, sorting, and concentration of individual target entities. This approach achieves the ultimate selectivity and sensitivity^12–15^.

In a recent development towards this aim, simulation studies proposed utilizing the mobility of biomolecules on flat 2D van der Waals material (vdWM) surfaces as a single-molecule detection and manipulation platform^16–28^. Several implementations have been proposed, including guiding DNA towards a nanopore by means of a graphene step edge^27^ and linearizing individual DNA and protein molecules by employing heterostructures of graphene/hexagonal boron nitride (hBN)/graphene^19,20,24^. The vdWMs exhibit a remarkable range of opto-electro-mechanical properties^29–32^, while also being biocompatible and robust in physiological environments^25,33–36^. Importantly, vdWMs can be seamlessly integrated into a single device and the large and atomically flat 2D surfaces of vdWMs provide ample space for the study of molecular dynamics and reaction processes, propelling the exploration of vdWMs as a new single-molecule platform in the fields of bioscience and biotechnology^37–41^.

Despite these promising prospects, to date, only few experimental studies have addressed the binding and diffusion of biomolecules on vdWMs at single-molecule resolution^42^. This pursuit has been hindered by the stringent requirements of the surface, which must possess three key properties: (i) optimal binding energy allowing mobility of biomolecules while ensuring prolonged localization, (ii) chemical stability and high cleanliness in aqueous environments, and (iii) no attributes such as fluorescence quenching or autofluorescence that could compromise optical measurement techniques including single-molecule fluorescence.

We identify hBN as a promising contender as it possesses outstanding chemical inertness and thermal stability^43–46^ and the wide energy bandgap of ∼6 eV precludes fluorescence quenching and autofluorescence^47–50^. Consequently, fluorophore-labelled molecules on a hBN surface could be detected even at the single molecule level, which was previously unattainable with graphene due to significant quenching effects^51–53^. In addition, hBN is also particularly well suited for studying the kinetics of DNA because the interatomic spacing of B and N atoms in the hexagonal 2D lattice closely matches, within 1.6 %, the C-C bond length of the hexagonal rings in nucleobases of DNA^43,54,55^. Thus, it is expected that the hydrophobic interaction due to these well-matched atomic distances leads to adsorption of DNA molecules. Moreover, it was predicted that graphene and hBN surfaces allow unimpeded lateral motion of molecules, enabling these to easily slide across the surface given the equivalent surface lattice sites^56^. This characteristic contrasts with more conventional surfaces, such as silicon nitride or silicon dioxide, which are known for their atomic roughness that significantly hinder lateral diffusion^57^. Despite these advantages, the detection of biomolecules and characterization of their dynamics on hBN surfaces has been limited, primarily due to technical challenges in imaging and a lack of complete understanding of the interactions between this solid-state surface and biomolecules.

Here, we present a comprehensive study of the binding and diffusion of DNA on hBN surfaces, using a combination of experimental and computational techniques. We find that single-stranded DNA (ssDNA) molecules bind non-specifically to the pristine, untreated hBN surface, resulting in their confinement within the 2D plane. The ssDNA molecules engage in diffusion on the surface of hBN, permitting us to explore the intricacies of their motion through a single-molecule tracking approach. Importantly, we reveal the role of several defect types on biomolecule motion and demonstrate that step edges and domain boundaries of hBN spatially constrain the motion of molecules. By engineering a narrow hBN ribbon structure, we further increase the confinement effect and clearly observe the motion of ssDNA molecules along the narrow channel, showcasing the potential application of hBN in nanofluidics and single-molecule transport.

## RESULTS AND DISCUSSION

### DNA Adsorption on hBN Surfaces

To investigate the interaction between DNA molecules and the hBN surface at single molecule resolution, we prepared 35 nt-long ssDNA molecules labelled with a Cy3 fluorophore. The hBN surface was prepared on a borosilicate coverslip via the mechanical exfoliation technique (Methods). Since the flakes on the coverslip were freshly cleaved, the topmost hBN surface was contaminant-free. Figure 1a illustrates the wide-field epifluorescence single-molecule microscope used to capture fluorescence images of a hBN flake immersed in a buffer solution carrying the ssDNA molecules.

**Figure 1.**
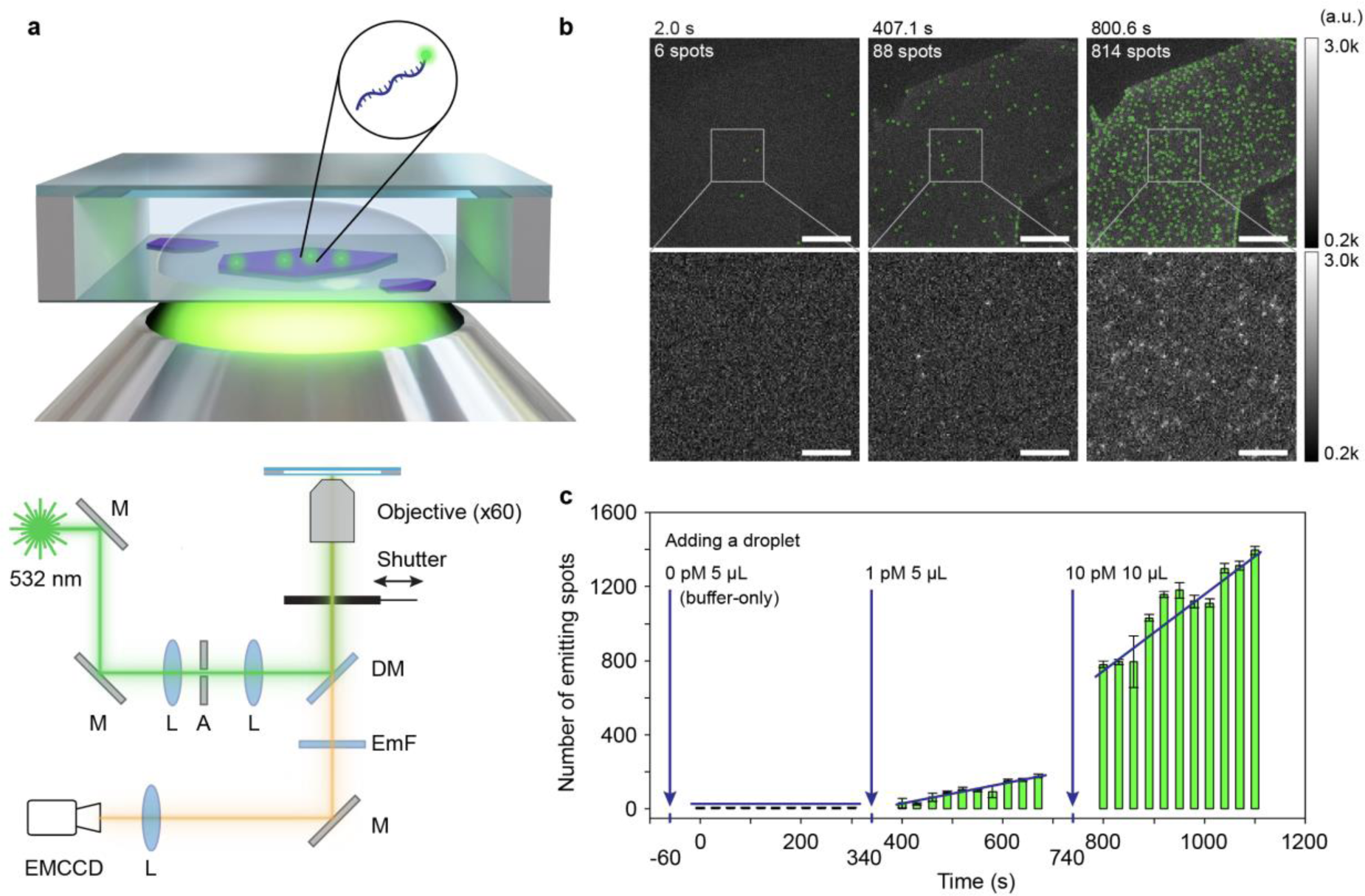
Single-molecule observation of ssDNA on the hBN surface. (a) A schematic representation of the sample chamber and measurement setup. Fluorescently labelled ssDNA molecules (green and blue) adsorbed onto the surface of an hBN flake (purple) are imaged using a wide-field epifluorescence microscopy setup. M, mirror; L, lens; A, aperture; DM, dichroic mirror; EmF, emission filter. (b) Fluorescence microscopy images of Cy3-labelled ssDNA molecules adsorbed on the hBN surface after injection of 0, 1, and 10 pM DNA concentrations (times correspond to Fig. 1c). ssDNA molecules are marked by a green circle. The bottom panels show the magnified view of the areas within the white boxes in the top panel. Both colour bars in the top and bottom panels display a range from 200 to 3000. The scale bars represent 20 μm (top) and 5 μm (bottom), respectively. (c) Temporal evolution of fluorescence emission spots. A 5 μL of droplet containing no DNA, 5 μL of 1 pM DNA and 10 μL of 10 pM DNA were added at –60 s, 340 s and 740 s, respectively (blue arrows).

We first measured the level of background signal of the pristine hBN surface by applying a 5 μl droplet of blank buffer solution. We observed ∼1 emitting spot per 1,000 μm^2^ on the hBN surface (Figures 1b and 1c). When we added a 5 μl droplet containing 1 pM ssDNA, we observed that the number of fluorescence spots increases at a constant rate. A higher rate was observed when an additional droplet of 10 pM ssDNA was added, providing evidence that the fluorescence spots correspond to ssDNA molecules. The ability to conduct studies at ultra-low concentrations, down to the low picomolar range, highlights the promise of hBN surfaces for single-molecule research.

### 2D Diffusion of DNA

During the single-molecule fluorescence measurements, it became apparent that the adhered ssDNA molecules moved over the surface, exhibiting substantial surface mobility, as predicted by previous molecular dynamics (MD) simulations^20,22–24,27,28^. The motion of the adsorbed ssDNA was notably faster when we used a shorter 7 nt (nucleotide)-long ssDNA instead of the long 35 nt ssDNA. For a quantitative analysis, we extracted trajectories of individual ssDNA molecules by using a single particle tracking algorithm (Figure 2a and Methods). Figure 2b shows the trajectories recorded over 400 s, and Figure 2c shows all the trajectories located within the square area. The 2D diffusion motion of ssDNA molecules on the hBN surface implies the surface binding energy of ssDNA is high enough to localize the molecule near the surface for a prolonged time but is also low enough to allow the molecules to diffuse along the surface.

**Figure 2.**
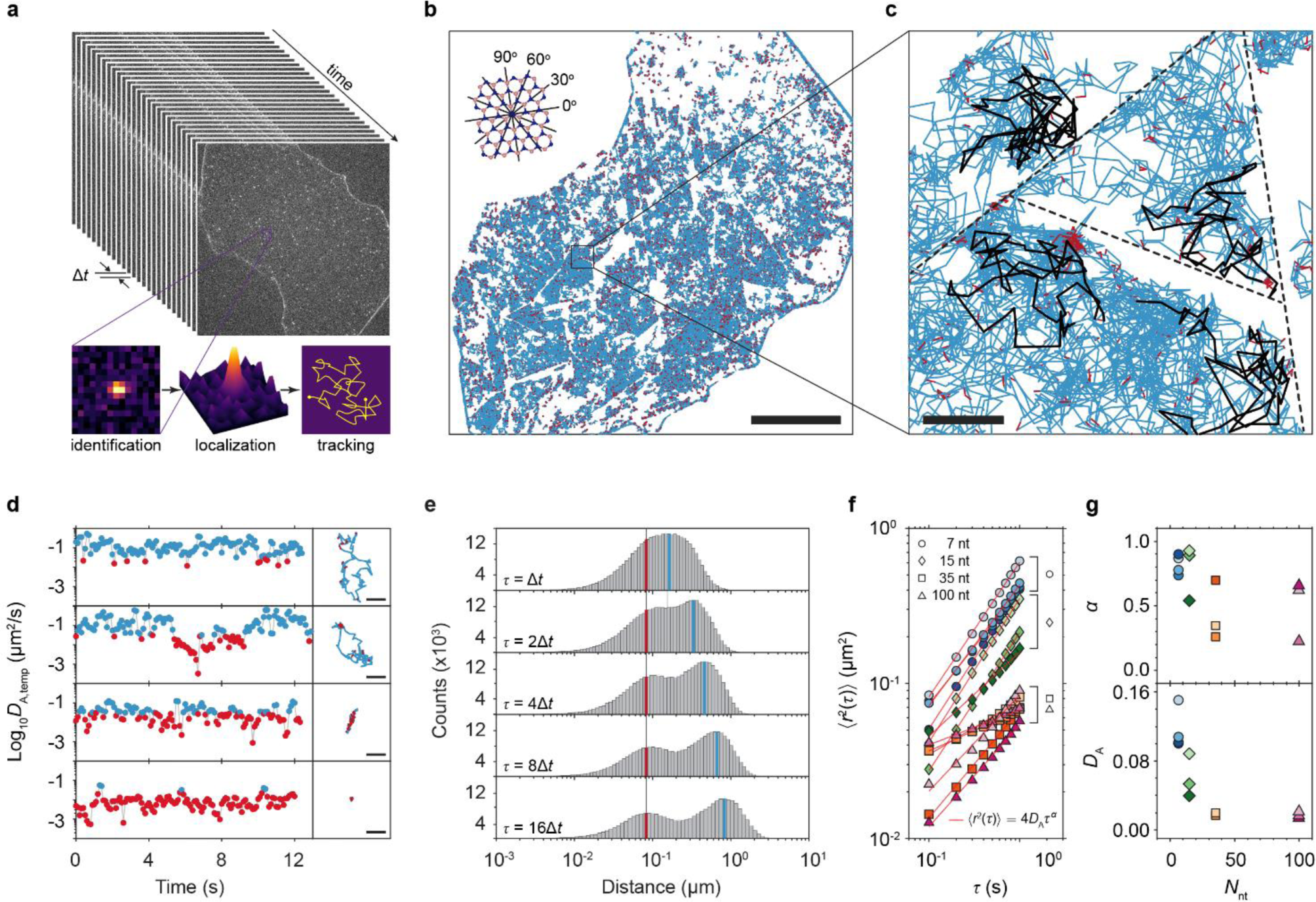
2D diffusion of ssDNA on hBN surfaces. (a) Schematic illustration of the single-molecule tracking analysis methodology. Identification of fluorescent spots in the fluorescence microscopy images were performed using differences of Gaussian (DoG) algorithm combined with a quadratic fitting scheme with sub-pixel resolution. The localized spot positions were then tracked across consecutive frames to establish molecule trajectories. (b) Trajectories of individual ssDNA molecules (7 nt) recorded over 400 s. Blue and red colours indicate mobile and stationary phases determined by temporal apparent diffusion coefficient (*D*_A,temp_) values (see Fig. d), respectively (see Methods for details). The lines in the top left corner denote the angles of the hBN lattice determined by the flake edges. The scale bar is 20 µm. (c) A zoom-in image from a square area in (b). The black trajectories are representative trajectories showing the confined movement in domain boundaries defined by step edges. The scale bar is 1 µm. (d) Apparent diffusion coefficient (*D_A_*) time traces of four representative molecules (left panels) with their 2D trajectories (right panels), with blue and red indicating the mobile and stationary states, respectively. (e) Jump distance distributions at various lag times *τ = n*Δ*t*, with *n* = 1, 2, 4, 8, 16 and the frame time Δ*t* = 0.1 s. The distributions show two distinct components, mobile (blue) and stationary (red). (f) Mean squared displacement ⟨*r*^2^(τ)⟩ for ssDNA molecules of different lengths plotted as a function of lag time τ. Data measured from different hBN surfaces are marked with different colours. (g) The apparent diffusion coefficient *D*_A_ and the diffusion exponent *α* in ⟨*r*^2^(τ)⟩ = 4*D*_*A*_τ^α^ are determined through linear fitting of the data in (f), as depicted by the red lines. The fill colours differentiate the data sets, which correspond to those in (f).

The trajectory maps revealed that there were boundaries on the hBN surface that ssDNA molecules were unable to traverse. Most boundaries were straight lines that run parallel to the lattice angles derived from the edge of the hBN flake (Figure 2b, top left). This confinement effect aligns well with recent MD simulations on ssDNA movement on graphene^27^. These simulations showed that the rate at which ssDNA moves across an atomic step is much lower than the rate at which it moves over a flat 2D plane since ssDNA molecules encounter resistance when they attempt to traverse the atomic step. This resistance exists not only at the up-steps but also at the down-steps, confining ssDNA effectively within the boundaries of a flat terrace on the hBN crystal.

Another significant aspect of Figure 2b is that the ssDNA molecules occasionally halt or considerably slow down over varying time periods, as highlighted in red. These instances of pauses and slow movements were identified based on whether the apparent temporal diffusion coefficient (*D*_A,temp_) fell below a certain threshold value of 0.031 µm^2^/s, equivalent to a single pixel area per second (see Methods). These events occur not only at step edges but also in the middle of a domain. This is more clearly illustrated in Figures. 2d and 2e, where *D*_A,temp_ and jump distances were derived from the trajectory data. Fig. 2d presents *D*_A,temp_ plotted over time as well as the corresponding trajectories, with blue and red colours indicating values that are above and below the threshold, respectively. The graphs from top to bottom display four molecules as representative examples, each demonstrating different types of motion: fast continuous diffusion, fast diffusion followed by stoppage, linear diffusion along a straight line and long-term anchoring at a single position. In Fig. 2e, the jump distances, the distance that a ssDNA molecule travels during a certain lag time *τ*, calculated for different values of *τ = n*Δ*t*, with *n* = 1, 2, 4, 8, 16 and Δ*t* = 0.1 s are shown. The histograms show two peaks, suggesting that the motion can be categorized into two states, slow (red line) and fast (blue line) diffusion. As the slow peak does not change its position with increasing lag times, we attribute it as a stationary state of molecules. We note that the jump distances measured from the stationary phase are ∼0.1 μm (red lines in Figure 2e), which is below the optical diffraction limit. Thus, the jump distances from the stationary phase probably merely reflect the uncertainty in the position determination from the single molecule images and do not necessarily correspond to motion of the molecule.

Further analysis using ssDNA strands with different lengths, specifically 7, 15, 35, and 100 nt, shows a decrease in both the apparent diffusion coefficient *D*_A_ and the diffusion exponent *α* values as the length of the DNA strand increases. Three datasets, all of which were measured from different hBN surfaces and are plotted separately in Fig. 2g, were collected per length of ssDNA. *D*_A_ and *α* were determined by fitting the average mean squared displacement (MSD) with the function ⟨*r*^2^(τ)⟩ = 4*D*_*A*_τ^α^, where ⟨*r*^2^(τ)⟩ and τ denote ensemble average of the displacement and the lag time, respectively (Figure 2f)^58^. Longer ssDNA molecules exhibit lower mobility *D*_A_ and more pronounced sub-diffusive behaviour, *i.e.* α < 1, indicating anomalous diffusion. Interestingly, *D*_A_ has a clear trend with length with no significant differences between the datasets, while α has a wide margin of error across the different datasets (Figure 2g). This indicates that DNA mobility is primarily affected by its length, while subdiffusivity is influenced not only by the DNA length but also by the state of the hBN surface, which is presumably the domain size determined by the step edges on the hBN surface.

### Computational Study of DNA Diffusion over hBN Surfaces

To elucidate the diffusion mechanism of DNA on the hBN surface, we performed MD simulations using the same parameters used in the experiments, i. e. the same buffer, salts, and the length of the ssDNA. As depicted in Figure 3, three types of hBN surfaces were simulated: a perfect planar surface (Figure 3a), a surface with a step edge (Figure 3b), and a surface containing 0.1 atomic % of B and N vacancies (Figure 3c).

**Figure 3.**
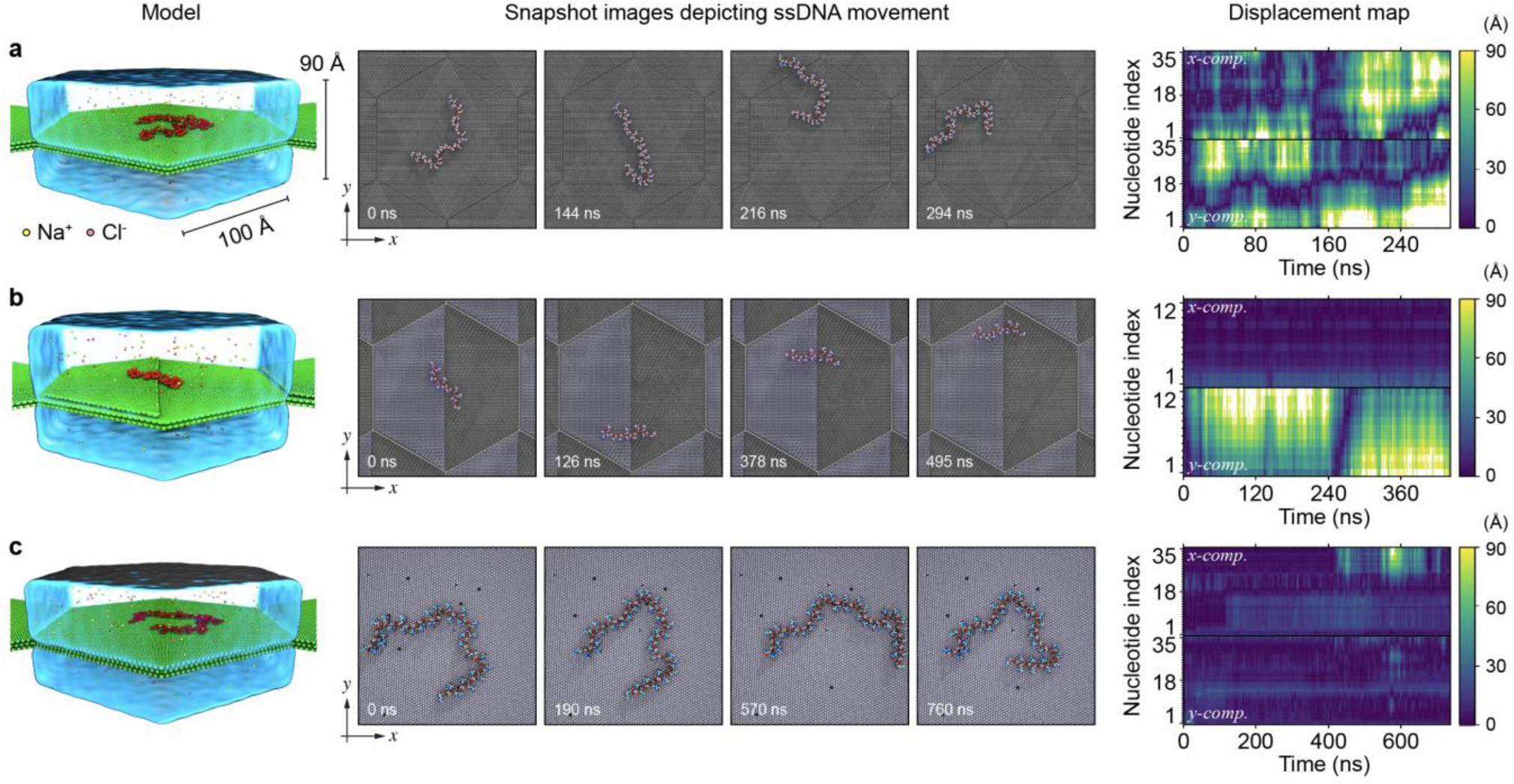
Molecular Dynamics (MD) simulation of ssDNA diffusion on hBN surfaces. Three distinct types of hBN surfaces were investigated: a perfectly flat surface (a), a surface with a single step edge (b), and a surface with atomic defects (c). The displacement maps on the last column show the displacement in x and y of each nucleotide as a function of time.

On the perfect planar hBN surface, the ssDNA molecule slid freely as visualized in the snapshot images in Figure 3a. The ssDNA molecules with lengths of 7, 15, and 35 nt exhibited normal 2D diffusion with diffusion coefficients of 749 μm^2^/s, 228 μm^2^/s, and 111 μm^2^/s, respectively. Notably, these values were significantly higher than the experimental values in Figure 2g by a factor 10^3^ – 10^4^. The MD simulation in Figure 3b also provides evidence supporting the possibility of linear movement of a ssDNA molecule along a step edge, similarly to the observation in the third row of Figure 2d. The displacement maps clearly illustrate the differences in DNA dynamics between a perfect planar surface and a surface with a step edge. On the perfect planar surface, high displacements occurred almost continuously in both the x and y directions. In contrast, on the surface with a step edge, displacements were not only reduced but also predominantly occurred along the y-axis where the step edge runs.

On hBN surfaces with 0.1% of atomic defects, the ssDNA molecule experienced a stationary state, as illustrated in Figure 3c. When the ssDNA molecule was bound to multiple atomic defects, it exhibited limited mobility within the group of defects. However, it was also observed that ssDNA was not permanently bound to the defects, and some segments of the molecule could escape from the defects, as depicted in the displacement map.

Our MD simulation revealed how ssDNA molecules interacted with hBN surface at nanosecond timescales, yet the large differences in the diffusion coefficients between the experiments and simulations remained to be explained. The MD simulations were limited to the (sub-)microsecond range, whereas the experimental timescale was several hundreds of seconds. To bridge this timescale gap, we combined the MD results with Monte Carlo (MC) simulations. We assumed that a freely diffusing molecule could be temporarily trapped on the surface when it encountered a defect site. As illustrated in Fig. 4a, the 2D diffusion was modelled with the following conditions: the DNA molecules diffuse on a finite square surface with area *L_d2_*, with a free-diffusion coefficient (*D*_0_) obtained from MD simulations of pristine hBN.

**Figure 4.**
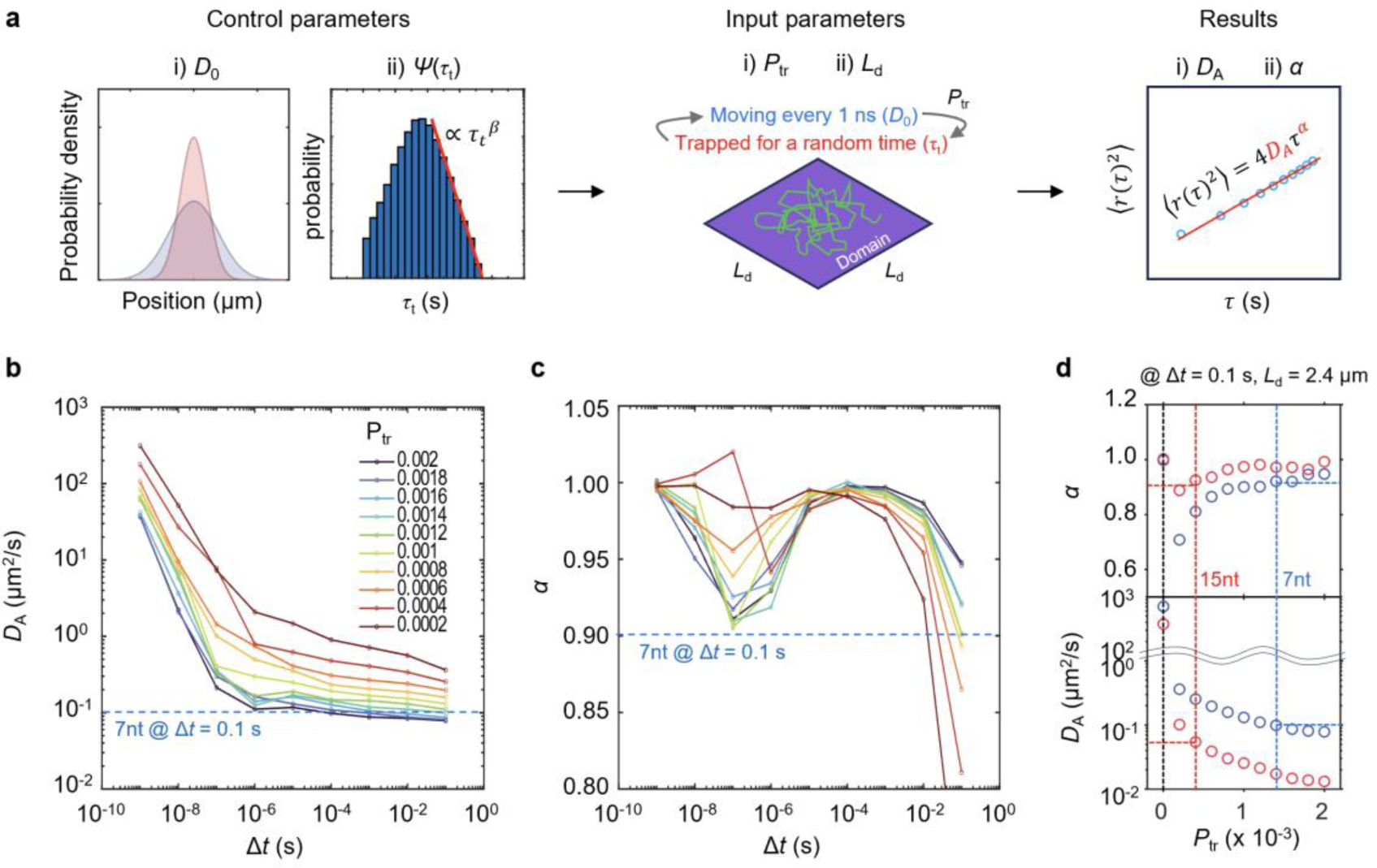
MC simulations of ssDNA diffusion on hBN surfaces. (a) Conceptual outline of the MC simulation model, which employs 100 particles that diffuse, with diffusion coefficient *D*_0_ determined from MD, in simulation steps of 1 ns with probabilities *P*_tr_ per step of encountering trapping sites. Trapped particles remain stationary for a random trap time *τ_t_* that is simulated using the probability distribution function *ψ*(*τ_t_*) before resuming movement. If a particle crosses the domain boundary *L*_d2_ based on its calculated displacement, its location is adjusted to be mirrored off the boundary, ensuring it remains within the domain boundary. Control parameters include a diffusion coefficient (*D*_0_) from MD simulations and a trap time distribution *ψ*(*τ_t_*) that is obtained from single-molecule fluorescence microscopy data. The domain length *L*_d_ is fixed at 2.4 µm and the trapping probability *P*_tr_ is varied in (b-d). By fitting the resulting MSDs, the effect of the domain size *L*_d_ and the trapping probability *P*_tr_ on the apparent diffusion coefficient *D*_A_ and the diffusion exponent *α* are determined. (b and c) Simulated variations in *D*_A_ and *α* for various *P*_tr_ are plotted as functions of frame time Δ*t*. (d) Influence of *P*_tr_ on *D*_A_ and *α* at set conditions of τ = 0.1 s and *L*_d_ = 2.4 µm, with blue and red lines representing experimental data for 7 nt and 15 nt ssDNA molecules. The corresponding *P*_tr_ values are 1.2 × 10^-3^ and 4 × 10^-4^ for 7 nt and 15 nt ssDNA molecules, respectively.

For 7 nt ssDNA, for which *D*_0_ = 749 µm^2^/s, we obtained the diffusion lifetime τ_d_ = 0.83 µs and the typical diffusion length *l_d_ =* 50 nm. For 15 nt ssDNA, we obtained τ_d_ = 2.5 µs and *l*_d_ = 47.7 nm using *D*_0_ *=* 228 μm^2^/s. The diffusion lengths of the two ssDNA with different lengths were similar to each other, suggesting that it is limited by surface defects. We can estimate the defect density to be approximately (1 cm/50 nm)^2^ cm^-2^ = 4 × 10^10^ cm^-2^, which is consistent with expected values for high-pressure grown hBN flakes^59,60^.

We conclude that the diffusion of ssDNA on the hBN surface has two phases. On a defect-free, flat hBN surface, ssDNA moves with a high diffusion coefficient of *e.g.* 749 µm^2^/s for 7 nt-long DNA. On a surface with atomic defects, the ssDNA can be intermittently trapped at defect locations. The trapping probability is estimated from MC simulations, from which we estimate a trap density of 4 × 10^10^ cm^-2^. The existence of these trapping sites can thus account for the experimentally observed apparent diffusion rates *D*_A_.

### Confinement of ssDNA molecules in a hBN nanochannel

In Fig. 2b, we discussed the confinement effects induced by step edges by limiting the motion of the ssDNA molecules within a single hBN terrace. This characteristic offers the opportunity to develop highly localized nanochannels that can guide DNA in a pseudo-one-dimensional manner. Notably, vdWMs like hBN possess an inherent property of cleaving along the crystal orientation, facilitating the preparation of clean ribbon-shaped crystalline surfaces. To demonstrate the feasibility of guiding molecules, we show in Figure 5 the motion of a 16-nt ssDNA on a narrow, elongated hBN nanochannel extending from a larger region of the crystal. Figure 5a represents a superposition of 3,500 images recorded during 350 s, where the letters c-f indicate the motion of a ssDNA molecule, shown in more detail in Figures. 5c-5f. As seen in Figure 5b, while the apparent diffusion coefficient (*D*_A_ = 0.039 μm^2^/s) value is comparable to that on a large 2D surface, we observe pronounced subdiffusivity (α = 0.64). Figures 5c-5f highlight various characteristic ssDNA movements on the hBN nanochannel, demonstrating ssDNA entering through an inlet (Figure 5c), moving linearly along the channel (Figure 5d), and navigating through y-junction branches (Figures 5e and 5f). These observations open the possibility of making a 2D fluidic device with hBN that can actively transport biomolecules on demand.

**Figure 5.**
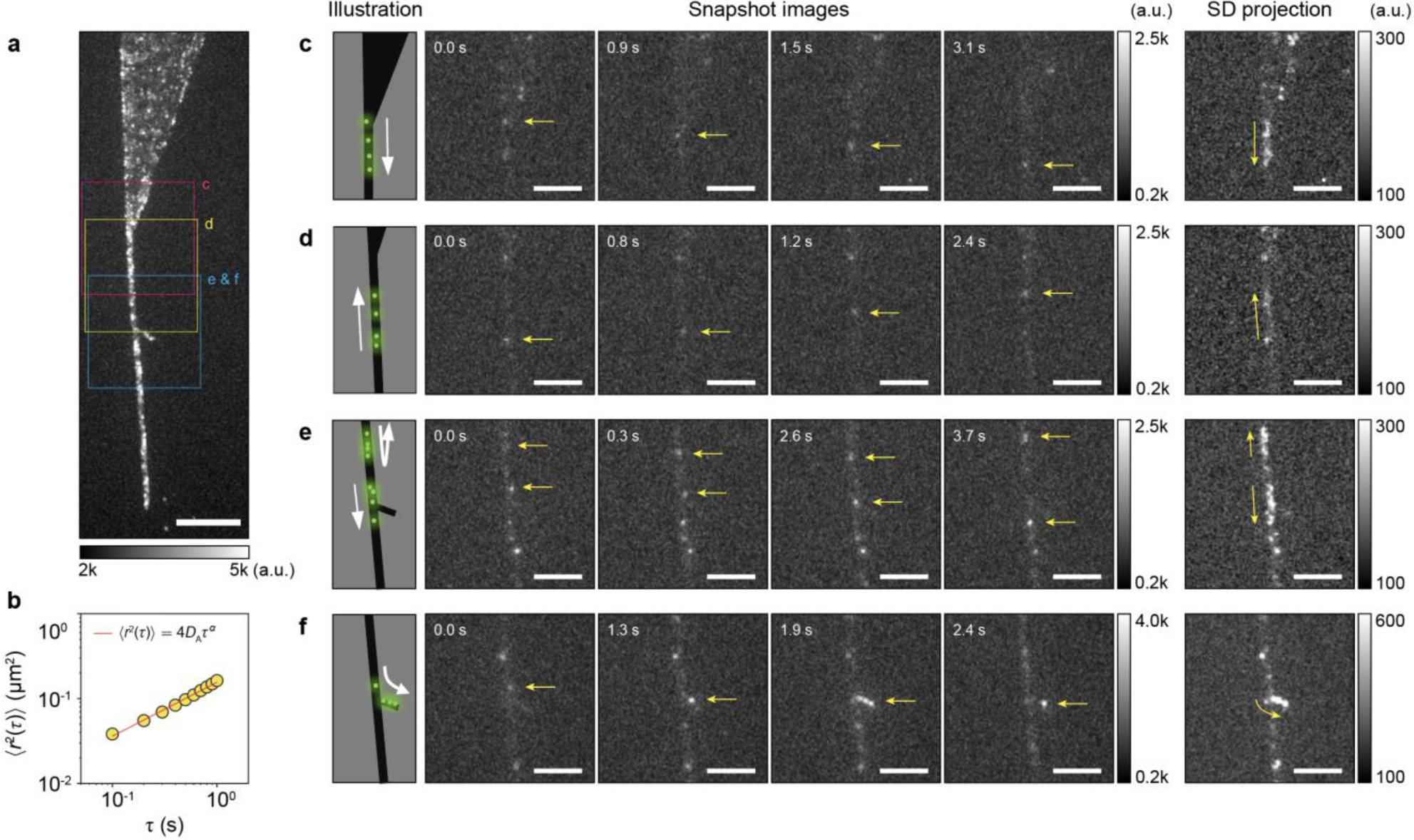
Surface diffusion of ssDNA molecules on a hBN nanochannel. (a) A time-based maximum projection of 3,500 epifluorescence microscopy images, showing 16 nt ssDNA molecules on a hBN surface. The hBN flake consists of three parts, a large reservoir (top), a narrow nanochannel (bottom), and a y-junction branch (middle). The scale bars are 10 μm. (b) The mean square displacement (MSD) as a function of the lag time, averaged over 292 trajectories. (c-f) Snapshots of ssDNA injection from the reservoir to the nanochannel (c), linear motion along the nanochannel (d), straight (e) and bending (f) movements at the y-junction. In the last column, the images show the standard deviation projection of the image stack over time, where the pixels with large intensity changes across the image stack are brighter. The presence of bright patches indicates the presence of mobile DNA molecules throughout the respective time periods (4.1, 2.4, 3.7, and 2.4 s for c, d, e, and f, respectively). The scale bar in (c), (d), (e), and (f) is 5 μm.

## CONCLUSIONS

We investigated the dynamics of DNA molecules on hBN surfaces using a multifaceted approach, yielding quantitative and mechanistic insights at single-molecule level. Our findings surpass the capabilities of traditional nano-imaging techniques. We showed that ssDNA molecules bind non-specifically to untreated hBN surfaces. Complementary computational studies with our single-molecule tracking results provided insights into the mechanisms of surface diffusion and confinement of ssDNA molecules as well as ssDNA-hBN interactions. Strikingly, our study revealed reduced mobility at step edges on hBN surfaces, resulting in the confinement of ssDNA molecules within the boundaries of the domains. These findings open avenues for constructing 2D nanofluidic devices featuring tightly constrained nanochannels, which will enable directional, one-dimensional motion of biomolecules, whether through surface diffusion or external stimuli. Additionally, engineering the defect density could be used to control diffusion rates, which holds significant potential for applications such as biomolecular sensors and bio-inspired electronic devices.

Atomically flat 2D surfaces have the potential to address fundamental biophysical questions and offer high-resolution sensing in confined environments. They eliminate the need of biochemical tethering strategies to bind molecules to a surface, nor do they rely on complex nanofabrication protocols to achieve confinement. However, achieving accurate and reliable biosensing depends on the precise manipulation and transport of biomolecules on these 2D surfaces, and these aspects continue to pose significant hurdles. For example, linearizing biomolecules like DNA and proteins allows for more accurate analysis and manipulation, but achieving this is not trivial. Additionally, implementing directional control can enable the selective and efficient transport of biomolecules to specific locations, opening a possibility for targeted drug delivery, high-throughput analysis, and accurate biosensing, yet this continues to present a significant hurdle. Our platform serves as a testbed offering high-throughput manipulation at single-molecule resolution including the high-resolution linearization of DNA. Furthermore, it can address these challenges through direct integration with active transport methods, such as dielectrophoresis and acoustofluidics, through which single-molecule directional transport can be tested with non-invasive and scalable device geometries.

To conclude, our study provides valuable insights into the binding, diffusion, and manipulation of DNA molecules on 2D surfaces, offering exciting prospects for the use of vdWMs as a solid-state platform for studying and controlling biomolecular dynamics in aqueous environments.

## METHODS

### 1) Preparation of the hBN surface

Highly crystalline hBN flakes were prepared by the mechanical exfoliation method^61^ on the borosilicate coverslip surface no more than 6 hours (typically 1-2 hours) before the measurement.

### 2) DNA and Buffers

Fluorescent dye (Cy3)-labelled single-stranded DNA (ssDNA) samples were purchased from Ella Biotech and used without further purification. Five ssDNA samples with different lengths were used in this study (7 nt, Cy3-CCTCCTC; 15 nt, AGATTTTTTTTTTTT-Cy3; 16 nt, Cy3-TTTTTTTTTTTTTTTT; 35 nt, AGATTTTTTTTTTTTTTTTTTTTTTTTTTTTTTTT-Cy3; 100 nt, AATGATACGGCGACCACCGAGATCTACACTCTTTCCCTACACGACGCTCTTCCGATCT ACGTATCACGAAAAAAAAATCXCGTATGCCGTCTTCTGCTTG), where X denotes amine-modified thymine base. The ssDNA samples were stored at 10 nM concentration in a buffer of 10 mM TrisHCl pH 8.0 and 50 mM NaCl. For each measurement, the ssDNA was diluted to a desired concentration immediately before applying it to the hBN surface. To avoid evaporation of the droplet on the coverslip, another coverslip was placed above the bottom coverslip with a ring or hollow square-shaped PDMS film (1 mm thickness) as a spacer (see Figure 1a).

### 3) Epifluorescence microscopy setup

To measure single-molecule fluorescence signals, a home-built epifluorescence microscope setup built around a commercial inverted microscopy body (Eclipse Ti2, Nikon) equipped with a high numerical aperture objective lens (60x, NA1.2, water immersion, Nikon) was used (Figure 1a). The beam size of a 532-nm laser CW diode-pumped solid-state laser (06-DPL 532nm, Cobalt) was enlarged by using a telescope and focused on the back focal plane of the objective lens for widefield excitation in epi-fluorescence mode. The Cy3 fluorescence signal was filtered by using a laser-blocking filter (ZET532/640m, Chroma Technology) and single-molecule fluorescence images were recorded by using an EMCCD (Ixon Ultra, Andor).

### 4) Single-molecule fluorescence measurement

For the single-molecule fluorescence measurement on hBN surfaces, we manually scanned for an hBN flake using a white light source. Once a flake was located within the field of view, we applied a 5 µL droplet of blank buffer (10 mM TrisHCl pH 8.0 and 50 mM NaCl) to the hBN flake and incubated it for 2 min. Following the incubation, we evaluated the level of background noise and contamination under 532 nm laser excitation. Afterwards, we added another 5 µL droplet of buffer solution (10 mM TrisHCl pH 8.0 and 50 mM NaCl) containing 1 pM ssDNA molecules to the existing droplet. After 2 min incubation in a dark room to allow the contents of the two droplets to mix, the fluorescence signals were recorded with a frame time of 100 ms. For the observations presented in Figure 1, the laser shutter was opened only for brief intervals, specifically 2-3 s every 30 s, to reduce fluorophore photobleaching. All the measurements were carried out at room temperature.

### 5) Single-molecule tracking

Single molecule images were analysed with the TrackMate plugin in the Fiji software^62,63^. Positions of individual fluorescent spots were determined by using differences of Gaussian algorithm combined with a quadratic fitting scheme with sub-pixel resolution^64^. To eliminate the interference of highly luminous spots near the flake boundary, the exterior of the boundary, beginning from its inner edge, was cropped out. Furthermore, a threshold for the sum intensity was applied to eliminate potential false-positive spots along the border, which might arise due to the stark contrast between the cleared and uncleared regions. Once the positions of individual molecules were ascertained, their trajectories across consecutive frames were determined using the Linear Assignment Problem mathematical framework^65^. The trajectory of a particle was determined by linking the particle’s position within consecutive frames if it is found in the next frame within 5 pixels (0.88 µm), as the probability of having a 2D displacement exceeding this value is less than 1% according to the analysis shown in Fig. 2e.

### 6) Diffusion analysis

The mean squared displacement (MSD) for a given lag time τ = *n*Δ*t* was calculated for each individual trajectory using 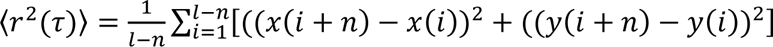 where *l* denotes the trajectory length, Δ*t* denotes the frame time, *n* denotes a positive integer^58,66^. *x*(*k*) and *y*(*k*) are the coordinates of the molecule at the frame index *k*. The MSD was calculated from the trajectories longer than 20 frames or more. The apparent diffusion coefficient *D*_A_ and the diffusion exponent *α* of the ensemble average of the MSD were determined by a linear fitting of the MSD plot using a 2D diffusion equation of 〈*r*^2^(τ)〉 = 4*D*_*A*_τ^α^.

At a specific frame index k with a time window T, the temporal MSD for the lag time *n*Δ*t* was calculated as 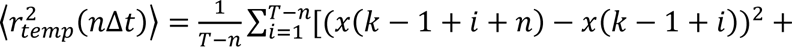 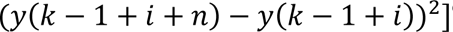^67^. From the temporal MSD, the temporal diffusion coefficient *D*_A,temp_ were derived: 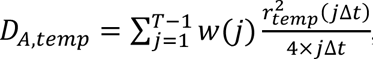, where the weighting factor 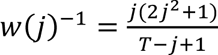 corresponds to the relative variance of 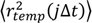^58^.

### 7) MD simulation

#### MD simulation methods

All MD simulations were performed using NAMD2.14^68^, the CHARMM36 parameter set^69^ for protein and DNA, TIP3P water model^70^ and along with the CUFIX corrections to ion-nucleic acid interactions^71^. The nonbonded interactions for B and N atoms in the hBN surface were set up according to previous work^72^. The hydrogen repartitioning scheme was used to achieve a 4 fs time-step^73^. Multiple time stepping was used^74^, with local and long-range interactions computed every 4 fs. All short-range nonbonded interactions were cut off starting at 1 nm and completely cut off by 1.2 nm. Long-range electrostatic interactions were evaluated using the particle-mesh Ewald method^75^ computed over a 0.1 nm spaced grid. SETTLE^76^ and RATTLE82^77^ algorithms were applied to constrain covalent bonds to hydrogen in water and in non-water molecules, respectively. The temperature was maintained at 295 K using a Lowe-Andersen thermostat^78^ with a cutoff radius of 2.7 Å for collisions and the rate of collisions at 50 ps*^−^*^1^. Energy minimization was carried out using the conjugate gradients method^79^. Initial equilibrations, *∼*20 ns, were performed in the constant pressure ensemble, using a Nose-Hoover Langevin piston with a period and decay of 2000 and 1000 fs, respectively^80^. Subsequently, all production simulations were performed in the constant volume ensemble. Throughout all simulations, the hBN surface atoms were fixed using harmonic position restraints with a scaling factor of 1 kcal mol^-1^ Å-2. Atomic coordinates were recorded every 9.6 picoseconds. Visualization and analysis were performed using VMD^81^ and MDAnalysis^82^.

#### Preparation of the MD simulation system

A hBN sheet consisting of 3 layers, was generated using VMD’s^81^ nanotube plugin. The 35 and 60 nt ssDNA were built using Avogadro^83^. Shorter sequences were generated by morphing residues from the 35 or 60 nt structures to match the desired ssDNA sequence using VMD’s PSFGEN plugin. The DNA molecule was then placed close to the hBN surface and submerged in a 50 mM NaCl solvent, using the solvate and autoionize plugins in VMD^81^. Required number of ions were determined by considering molality (mol kg*^−^*^1^). All systems were simulated with hexagonal periodic boundary conditions. Atoms on the boundary were covalently bonded across the periodic lattice, to create an effectively infinite hBN surface. For the hBN surfaces, three distinct types were simulated: i) a perfectly flat surface, ii) a surface with a single step edge, and iii) a surface with atomic defects. The hBN surface with a single step edge was constructed by removing one half of the atoms on the topmost hBN layer. The partial charge on atoms present on the defect boundary was set zero, to avoid unphysical pinning of nucleotides at the boundary. The hBN surface with atomic defects were created by random removal of B and N atoms on the topmost hBN surface. Here, the stoichiometry between B and N atoms, and hence the charge, was preserved during the removal of atoms.

#### Summary of MD simulation systems

In total, we simulated ten independent systems, each differing by the ssDNA sequence, the nucleotide count, and the presence of defects on the hBN surface. Four systems contained a poly(dAdT) DNA homopolymer of comprising of either 7, 15, 35 or 64 nucleotides were simulated on a perfectly flat hBN surface, i.e. devoid of any defects for ∼300 ns each. These systems contained approximately 250k, 250k, 390k and 700k atoms respectively, and each of these were simulated for *∼*300 ns. Another 250k-atom system contained a poly(dAdT)_12_ strand and an hBN surface having a single step edge and was simulated for *∼*500 ns. Another three systems contained a poly(dAdT)_35_ strand and an hBN surface with atomic defects produced by randomly removing 1, 0.2 or 0.1% of both B and N atoms. Finally, two separate ∼250k-atom systems contained either a poly(dA)_25_ or poly(dT)_25_ strand and were each simulated in the presence of a smooth hBN surface for ∼300 ns.

## ACKNOWLEDGMENTS

CJ was supported by Human Frontier Science Program (RGP00026/2019), NWO (Vici), the Basic Research Laboratory Program (NRF-2023R1A2C2004745), and Frontier 10-10 (Ewha Womans University). KC and AA were supported by the US National Science Foundation (DMR-1827346) and the Human Frontier Science Program (RGP0047/2020). The supercomputer time was provided through ACESSS allocation grant MCA05S028 (A.A.) and the Leadership Resource Allocation MCB20012 on Frontera of the Texas Advanced Computing Centre (A.A.). SC was supported by the ERC starting grant (SIMPHONICS, No. 101041486) and a Delft Technology Fellowship. DHS was supported by the KIND fellowship program at the Kavli Institute of Nanoscience Delft and the ERC starting grant (SIMPHONICS, No. 101041486). SK was supported by Brain Pool program funded by the Ministry of Science and ICT through the National Research Foundation of Korea (RS-2023-00261876).

